# Hydrogen Sulfide Suppresses Keratinocyte Migration in Scratch Assay

**DOI:** 10.1101/2025.08.13.670126

**Authors:** Alan Chen, Vijay Krishna

**Author notes:** Vijay Krishna; 9500 Euclid Avenue, Cleveland, Ohio 44195.

## Abstract

Cell migration is crucial for maintaining physiological functions in the body and is driven by mechanical, electrical, or chemical stimuli. Hydrogen sulfide (H_2_S), a gaseous signaling molecule, has been shown to regulate cellular migration in a cell type and context-dependent manner. Keratinocytes are the predominant cell-type in the epidermis and drive re-epithelialization during wound healing. However, the effect of H_2_S on keratinocyte migration remains incompletely defined. Here, we demonstrate that H_2_S suppresses keratinocyte migration in a serum-restricted *in vitro* scratch assay. Intracellular H_2_S levels were modulated using fast-releasing donors H_2_S donors (NaHS or Na_2_S) to achieve supraphysiological levels or by siRNA knockdown of endogenous H_2_S-producing enzymes *cystathionine-γ-lyase* (CGL) and *cystathionine-β-synthase* (CBS) to obtain infraphysiological levels. Treatment with NaHS or Na_2_S (10 µM–5 mM, 24 h) reduced gap closure in a dose-dependent fashion, with negligible effects at ≤ 100 µM and progressive suppression at higher concentrations. Conversely, siRNA knockdown of CGL or CBS boosted keratinocyte migration by 30% versus scrambled siRNA control. Supplementation with a slow-release H₂S donor rescued the siCGL phenotype, restoring migration to control levels. Across experiments, intracellular H₂S level is negatively associated with keratinocyte migration. These findings suggest that H₂S can restrain keratinocyte motility under serum-restricted conditions, and that reports of accelerated wound closure with H₂S donors *in vivo* may reflect effects on other cell types and/or inflammatory pathways rather than direct enhancement of keratinocyte migration.

**Significance Statement:** Dysfunctional wound healing affects millions worldwide. While H₂S donors can accelerate wound closure in animal models, the cell-type–specific effects remain unclear. We show that, H₂S suppresses keratinocyte migration implicating non-keratinocyte mechanisms (e.g., endothelial, immune, fibroblast) in the *in vivo* benefit of H₂S donors and refining where in the healing cascade H₂S is most likely to help.

## Introduction

In the United States, dysfunctional wound repair affects more than 10 million people, annually, where approximately 300,000 patients are hospitalized (1). Chronic conditions such as diabetes and cardiovascular disease increase risk and delay wound closure, underscoring the need for targeted pro-healing strategies (2–4). Wound healing progresses through inflammatory, proliferative, and remodeling phases, each involving coordinated migration of platelets, keratinocytes, immune, microvascular, and fibroblast populations. During the remodeling phase, cells tend to stop proliferating and generally migrate (5, 6). Cells migrate in response to various stimuli (i.e., mechanical, electrical, chemical) via changes in their intracellular cytoskeleton structure, cell-cell interactions (e.g., cadherins, gap junctions), and/or cell-matrix interactions (e.g., integrins) (7).

Hydrogen sulfide (H_2_S) is one such chemical stimuli that has been shown to regulate cellular migration (8). Endogenously, H_2_S in mammals is produced predominantly by enzymes of the transsulfuration pathway, such as *cystathionine-β-synthase* (CBS) and *cystathionine-γ-lyase* (CGL), but can also be produced non-enzymatically via a combination of free or heme-bound iron and vitamin B6 (8, 9). Previous studies have demonstrated that the effect of H_2_S on migration is cell-type and environment dependent. For example, studies utilizing endothelial and cancer cells have reported that H_2_S promotes cell migration *in vitro*. By contrast, studies that uses other non-cancerous cell types (e.g., smooth muscle and immune cells) have reported that H_2_S suppresses cell migration *in vitro* (10, 11). H_2_S have been found to affect cell migration through the Akt pathway and mitochondrial energetics (12).

In terms of skin wound healing, exogenous H_2_S applications have been found to promote wound closure in murine, rat, and non-human primate models (5, 10, 13). Further, in late stage wound repair (where migration predominates), CGL expressions have been observed to be elevated in the skin *in vivo*, implying a role of endogenous H_2_S pathways in wound healing (5). Several *in vitro* studies have demonstrated evidence in line with these pre-clinical observations by correlating H_2_S’s therapeutic effect with cell proliferation and differentiation during the early stages of wound repair in endothelial and fibroblast cells (10, 11). In addition, H_2_S have also been found to promote migration that is common during the late stages of wound repair in these cell types (endothelial and fibroblast cells) (10, 11). However, despite being the most predominant cell type in the skin during late-stage wound repair where migration predominates (14), the effect of H_2_S on keratinocyte migration is lacking.

In this study, we have investigated the role of H_2_S on keratinocyte migration in an *in vitro* scratch assay. H_2_S levels can be modulated by use of either exogenous chemical donors (e.g., sodium salts and pro-drugs) or through its endogenous production from sulfur amino acid catabolism (10–12).

## Results and Discussion

### Cell Migration Image Analysis

Cell migration was evaluated using an *in vitro* scratch assay (15, 16). A gap area was formed by scratching a confluent cell monolayer grown on polystyrene plastic with a sterile 10p pipette tip and straight edge (**Figure 1, Top**). Cells were scratched in the same direction and position in the biosafety cabinet to minimize any artifacts of scratching (i.e., pressure, gap length variations) across replicates (15). To minimize effect of proliferation, cell migration assays were conducted under serum starvation (1% FBS) conditions (6). Cell migration was observed with an environmentally controlled (37°C, 5% CO2) time-lapse phase contrast microscope that took snapshots of the migrating cell front over a 24-hour period (**Figure 1, Middle; Supporting Video 1**). The gap closure was quantified by measuring the change in gap area over time using the open-source CellProfiler Cell Imaging Software (17) (**Figure 1, Bottom**). Gap area, as opposed to gap width/length, was used as the final endpoint for migration as it better represents the multidirectional movement of cells (15, 18). The image processing parameters for CellProfiler were set for analyzing all images and details are provided in Materials and Methods section (16, 19) (**Supporting Figure 1**). The algorithm was validated by comparing the output images and gap areas from CellProfiler to manual freeform area measurements in ImageJ (19), at 0, 6, 12, 18, and 24 hours (**Figure 1, Bottom**). There was no significant difference between the gap areas determined from CellProfiler and ImageJ, thus validating the algorithm (two-tailed Student’s t-test, α = 0.05).

**Figure 1:**
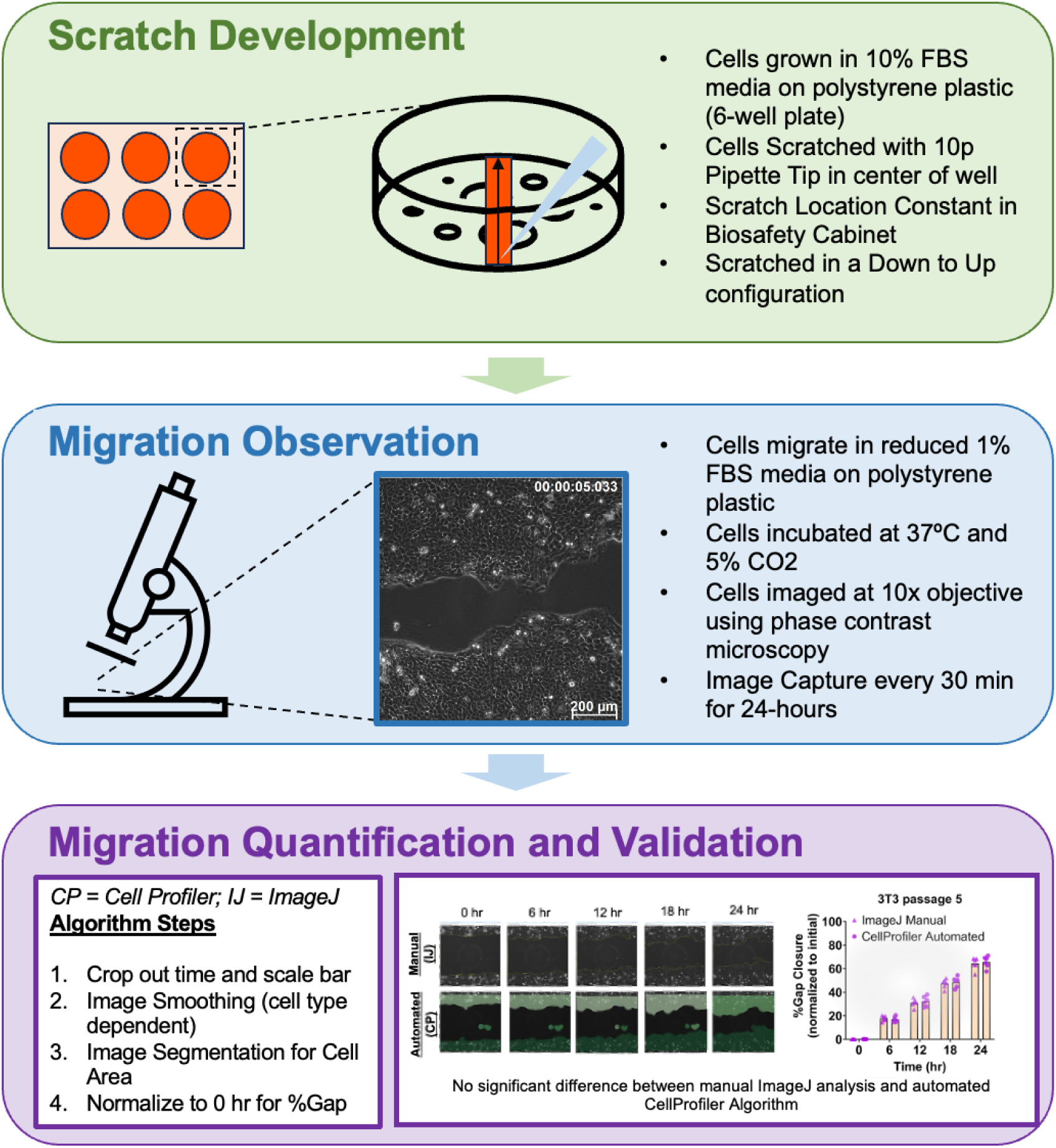
Development of scratch assay consisting of scratch development (**top**), migration observation (**middle**), and migration quantification (**bottom**). A customized set of parameters in CellProfiler was used to quantify changes in gap area as an indication of migration. The automated quantification was validated with manual analysis using ImageJ at selected time points 0, 6, 12, 18, and 24 hr (n = 6; mean ± SD) (Student’s t-test).

### Exogenous H2S suppresses keratinocyte migration

To evaluate the effect of exogenous H_2_S on HaCaT keratinocyte migration, we used NaHS and Na_2_S as fast-releasing H_2_S donors prepared fresh in buffered media (**Supporting Figure 2**). After 24 hours, a negative correlation between H_2_S concentration and HaCaT keratinocyte migration (represented as % Gap Area) was observed (**Figure 2**). Statistically, low concentrations (< 100 µM) of H_2_S did not contribute significantly to migration compared to control while higher concentrations seem to suppress HaCaT migration in a dose dependent manner (One-Way ANOVA) (**Figure 2**).

**Figure 2:**
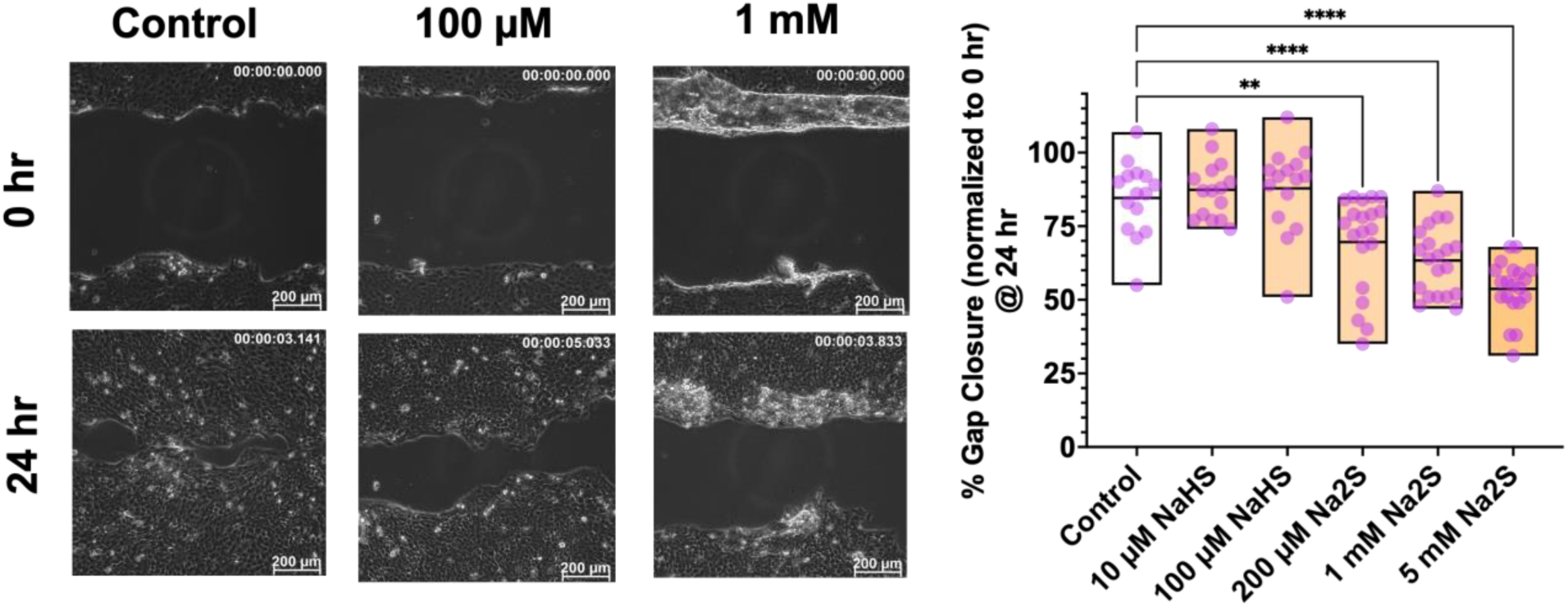
Contribution of exogenous H_2_S Donors on HaCaT keratinocyte (passage 60 ± 2) migration. Effect of exogenous H_2_S was investigated using fast-acting donors NaHS and Na_2_S for low and high concentrations, respectively. Snapshot images of HaCaT migration with normal or added exogenous H_2_S (100 µM and 1 mM) are shown for 0 and 24 hr. Analysis of keratinocyte migration with exogenous H_2_S treatment at 24 hours is quantified using % Gap Closure (normalized to initial time point) (n = 15-20).

Previous studies have also observed that H_2_S either suppressed or didn’t affect cell migration in various non-cancerous and non-endothelial cell-types (8). The effect of serum starvation is critically important in the analysis of cell migration as it could influence the degree of cell proliferation (11). In a study where immortalized keratinocytes (HaCaT) were treated with 100 µM H2S donor in a scratch assay under serum-starved conditions, it was observed that the treatment had no effect on proliferation, viability, adhesion or migration of HaCaT cells, which is in line with observations at 100 µM in **Figure 2** (11). Only after an inflammatory condition was introduced to the same system (e.g., methylglyoxal injury) was a therapeutic effect of H_2_S observed in HaCaT cells (11), suggesting a potential contribution of the immune system that could explain the observations from various *in vivo* models (5). Further, it is generally accepted that the H2S response curve for most biological responses is an inverted bell shape, meaning that at a certain concentration, a therapeutic effect should persist, but results from **Figure 2** does not show this expected response (8). Only one study investigating the effect of 50 µM NaHS on HaCaT migration under serum-starved conditions has demonstrated a slight but statistically significant advantage of H_2_S compared to control (10). This suggests that there could be a potential benefit of H_2_S on HaCaT migration between the concentrations of 10 and 100 µM in **Figure 2**.

### Knockdown of H2S-producing enzyme accelerates keratinocyte migration

The effect of endogenous H_2_S on HaCaT migration was investigated by using siRNA to knockdown H_2_S-producing enzymes CGL or CBS. A scrambled set of siRNA was used as control for these experiments. Similar to the exogenous H_2_S experiments, HaCaT cells were allowed to proliferate for 72 hours before initiating the scratch assay in 1% reduced serum media. During the initial 72-hour phase, cells were first reverse transfected for 48 hours with siRNA-lipofectamine before being forward transfected another 24 hours to ensure efficient knockdown of target enzymes (CGL or CBS). The siRNA knockdown of H_2_S-producing enzyme showed 30% reduction in endogenous H_2_S levels (as measured via p3 fluorescence (20)) for both enzymes (CGL or CBS) when compared to the control (**Supporting Figure 4**). Low magnitude of H_2_S reduction could be attributed H_2_S production from non-targeted H2S-producting enzymes (e.g., CBS for siCGL, CGL for siCBS, and 3-MST) (13). Importantly, knockdown of H_2_S-producing enzymes increased HaCaT migration by approximately 30% (**Figure 3**).

**Figure 3:**
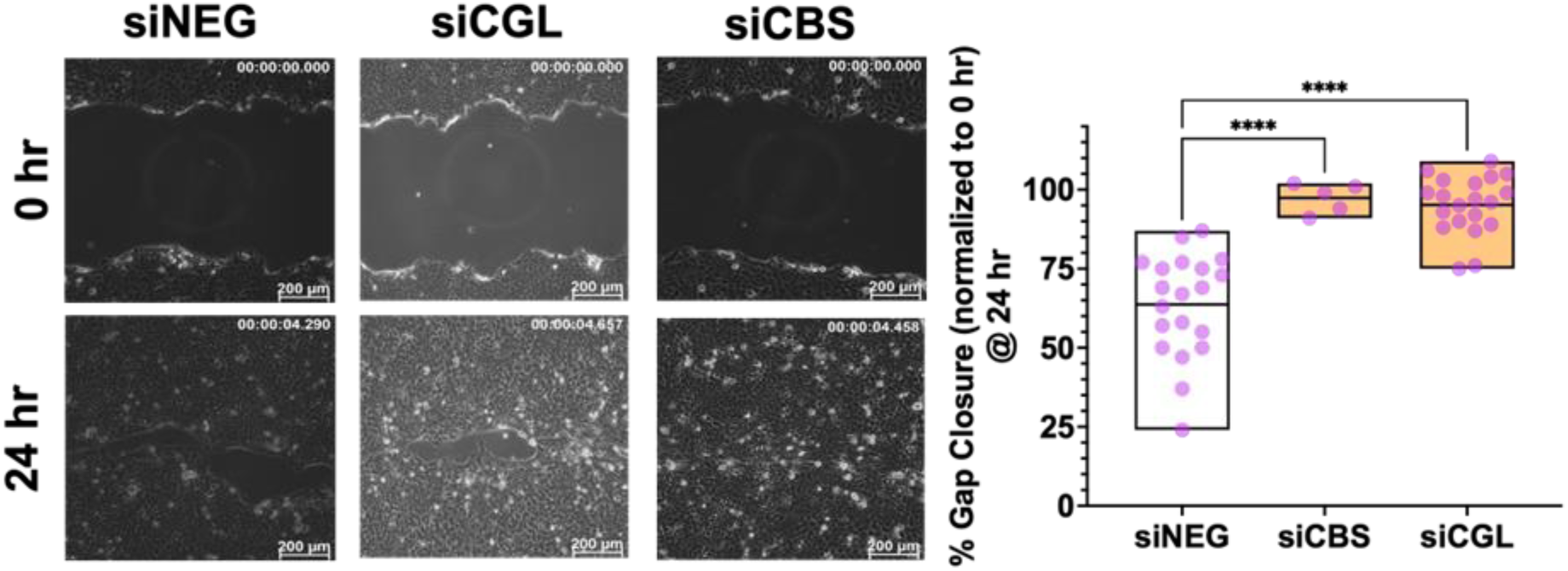
Contribution of endogenous H_2_S on HaCaT keratinocyte (passage 60 ± 2) migration. Effect of endogenous H_2_S was investigated using siRNA for silencing CGL or CBS. Snapshot images of HaCaT cell migration with normal or limited endogenous H_2_S are shown for 0 and 24 hr. Analysis of keratinocyte migration at 24 hours was quantified using % Gap Closure (normalized to initial time point) (n = 5-20).

The effect of H_2_S observed in both **Figures 2** and **3** have also been reported *in vitro* scratch assays by various other researchers. For example, Yang et al., reported that exogenous NaHS suppress migration in smooth muscle cells, whereas reduction in H_2_S generating capacity in these cells via CGL knockout demonstrated the reverse effect – an acceleration in cell migration (21). In contrast, the study that investigated the role of H_2_S on endothelial cell migration in angiogenesis demonstrated that addition of 60 µM NaHS stimulated cell migration while addition of CGL siRNA reversed that effect (22). This implies that H_2_S has an overall suppressive role in non-endothelial cells (8).

### H2S donor rescues the knockdown of H_2_S-producing enzyme

To validate the overarching hypothesis that H_2_S suppresses HaCaT keratinocyte migration, next set of experiments utilized a proprietary H_2_S donor to rescue the diminished levels of H_2_S in siCGL treated cells. The release of H_2_S from proprietary donor can be observed visually as brown bismuth sulphide spots on bismuth acetate infused paper (Na_2_S was used as positive control) (**Figure 4A**). Addition of the proprietary donor induced a dose-dependent suppression of keratinocyte migration that reversed the accelerated migration observed in the siCGL back to control (siNEG) levels, thus validating the initial hypothesis (**Figure 4B**). Based on the results and discussion in the manuscript, H_2_S clearly plays a suppressive role in the migration of HaCaT keratinocytes.

**Figure 4:**
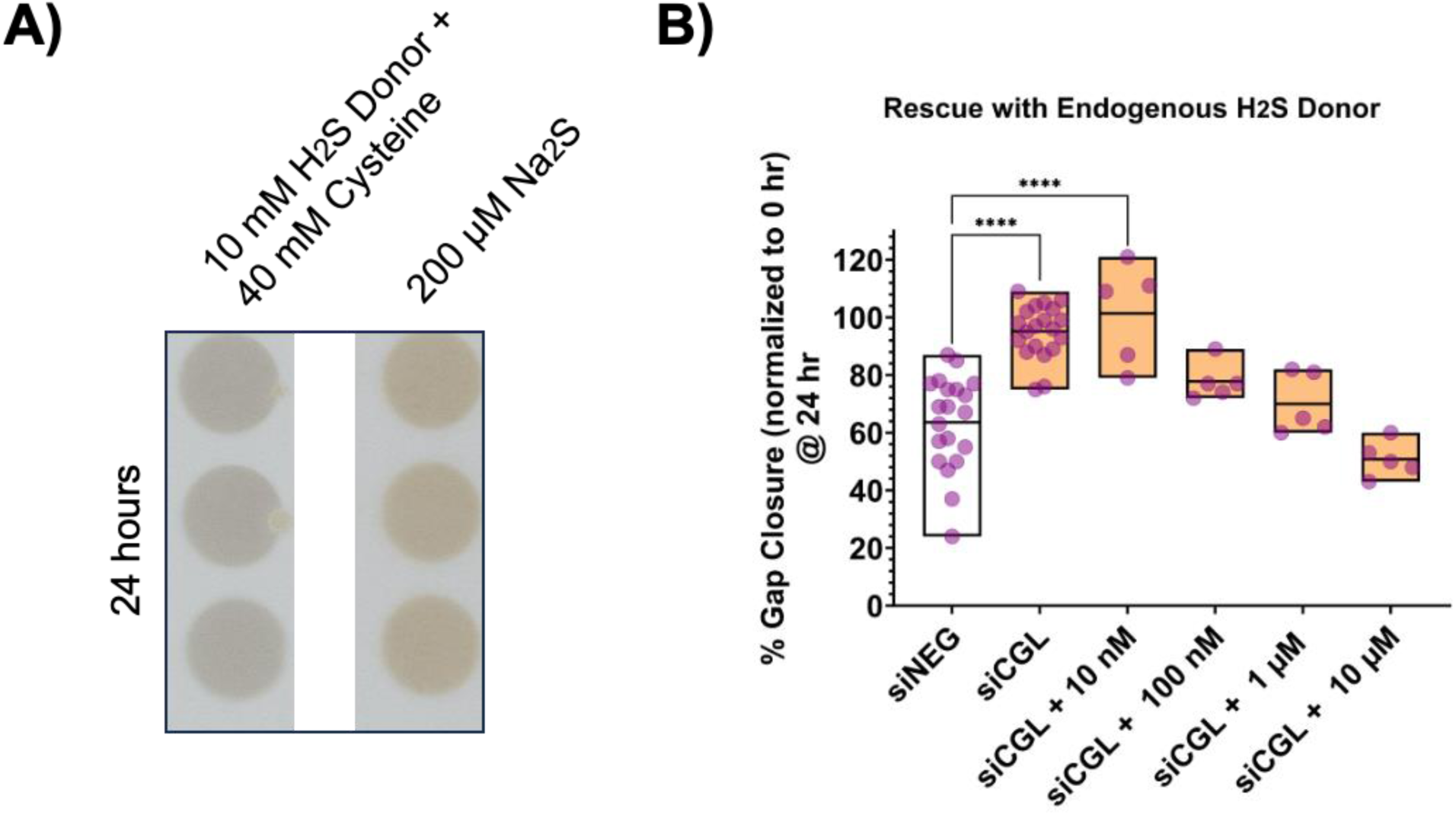
(A) Production of H_2_S by 10 mM proprietary donor after 24-hours. H_2_S was detected on bismuth infused colorimetric paper and compared to positive control (200 µM Na2S). **(B)** Rescue experiment using a proprietary H_2_S donor on siRNA treated HaCaT keratinocyte (passage 60 ± 2) migration. H_2_S supplementation reverses the effect of siCGL. Analysis of keratinocyte migration at 24 hours was quantified using % Gap Closure (normalized to initial time point) (n = 5-20).

### Limitations of Study

The observations reported in this study were generated with immortalized keratinocytes grown under static conditions. Immortalized cells were used due to their simplicity, accessibility and affordability (23), however, they have been found to be prone to oncogenic and genetic drifts, which can confound results (24, 25). Several studies have also directly or indirectly implied the effect of immune system markers on cell migration (11, 26). Hoyle et al., have shown that the time of the day for scratch assay can inadvertently affect cell migration in both primary culture and HaCaT cells (27). Our study did not control the time of scratch initiation. Further, cell behavior *in vivo* is also affected by a multitude of factors such as fluid flow, pressure, and multi-cell interactions, which are lacking in this simple monoculture model.

### Future Direction

The overall goal of this study was to determine the effect of H_2_S on keratinocyte migration. Our data, along with observations for other cells in literature, support the suppressive role of H_2_S on cell migration. This conclusion implies that H_2_S’s therapeutic effect observed in vivo may not be dependent on keratinocyte migration but rather other areas of the wound healing cascade. Further studies would need to be conducted to delineate the effect of H_2_S on 3-dimensional cell culture containing keratinocytes and other cell types to better understand the role H_2_S on wound healing.

## Materials and Methods

### Chemicals and Reagents

NaHS hydrate (#161527), Na_2_S (#407410), and p3 dye (#5.34329) were acquired from Sigma Aldrich (St Louis, MO). Pro-Long Diamond Anti-Fade Mountant with DAPI (#P36971), LEICA Type F Immersion Liquid (#11513859) and 0.4% Trypan Blue (#15250-061) were acquired from ThermoFisher (Waltham, MA). Corning 6-well tissue culture treated plates (#3526), ibidi 8-well glass chamber slides with silicone inserts (#80841), T-75 cell culture flasks (#12-566-85), Eppendorf 10µL (# 05-414-048) and 1 mL pipette tips (# 05-413-996) were acquired from Fisher Scientific (Hampton, NH). 200 µL pipette tips (# 4-1059-225-000) were acquired from LabCon (Petaluma, CA). Cell culture DMEM (CaissonLabs, #DMP08), 1x PBS (CaissonsLabs, #PBP01), 0.25% Trypsin and 0.53 mM EDTA were acquired from the Lerner Research Institute Cell and Media Core.

### Cell Culture

NIH-3T3 immortalized embryonic fibroblasts were acquired from ATCC (NIH/3T3 (ATCC CRL-1658)). HaCaT immortalized keratinocytes were a gift from Dr. Nora Singer at Cleveland Metrohealth. Cells were grown on T-75 flasks in 10% FBS, 1% P/S DMEM media with 10 mL media being replenished every other day in a 37°C, 5% CO_2_ incubator (ThermoFisher Heracell™ VIOS 160i). Cells were detached from T-75 flask with 5 mL 0.25% trypsin and 0.53 mM EDTA. HaCaT cells were used from passages 58 to 62. 3T3-NIH cells were used from passage 1-5. When ready for experiments, cells were counted using the Countess 3 FL Automated Cell Counter (ThermoFisher) with 1:1 cell suspension to trypan blue ratios (total volume = 20 µL).

### siRNA Complexing

Ambion *Silencer* siRNA for CGL (cystathionine-γ-lyase) (siRNA # = s99002; Sense CGAUUACACCACAAACCAAtt) was used to knock down CGL from cells. Ambion *Silencer* siRNA for CBS (cystathionine-β-synthase) (Part # = s63476; Sense GCAAAGUCCUCUACAAGCAtt) was used to knock down CBS from cells. Ambion *Silencer* Negative Control 1 (Part # =4390843) was used as the scrambled RNA control for RNA silencing experiments. siRNA was complexed and delivered to cells using lipofectamine RNAiMax lipofection. Briefly, 6.25 µL lipofectamine RNAiMax in 250 µL DMEM media without serum and antibiotics, was added to 17.5 µL of 28 pmol siRNA in 250 µL DMEM without serum and antibiotics for 10 minutes at room temperature. The final concentration of siRNA in complexing media was 100 nM with a total volume of 500 µL. Volumes indicated are for cellular transfection in a 6-well corning multi-well tissue culture treated plates with a total media volume of 3 mL. Protocol was adapted from Invitrogen’s protocol for transfecting siRNA-RNAiMax into NIH-3T3 cells (28). For transfection performed with other media volume, amount of siRNA and components used were scaled accordingly.

### siRNA Transfection

HaCaT cells was first reverse transfected with siRNA for 48 hours and then subsequently forward transfected for an additional 24 hours (total transfection time = 72 hours). In tissue culture coated 6-well plates, siRNA was first complexed with lipofectamine in DMEM media without serum and antibiotics for 10 minutes at room temperature. Subsequently, cells in DMEM media (10% FBS, no antibiotics) were seeded at a density of 300,000 cells/well and diluted to a total volume of 3 mL. After 48 hours, 500 µL of media was removed and replaced with 500 µL of siRNA-RNAiMax complex to initiate forward transfection. For transfection performed with other media volume, amount of siRNA and components used were scaled accordingly.

### Detection of H_2_S with p3 fluorescent probe

For p3 probe assay, 50,000 cells were seeded into each well of an ibidi glass 8-chamber well slide with silicone inserts. siRNA transfection was conducted accordingly, and amounts were scaled to the total volume used (500 µL). After siRNA transfection, cells were immediately rinsed once with 1xPBS and then incubated with 500 µL of 20 µM p3 probe (p3 probe was added via a five-fold dilution) for 10 minutes to detect endogenous H_2_S levels. Next, cells were rinsed once with 500 µL 1x PBS and then fixed with 500 µL 4% paraformaldehyde (diluted in 1x PBS) for 10 minutes. After fixation, cells were washed for 5 minutes one time with 1x PBS buffer and then silicone inserts were removed from the slide. The slide was gently tapped on its sides to remove extra water that remained on the slide. 2 drops of 50 µL of Pro-Long Diamond Anti-Fade Mountant with DAPI was then added to a cover slip, which was immediately covered onto the cells. Mounting media was allowed to cure, in darkness, overnight at 4°C until hardened. P3 probe was then detected using a Leica TCS-SP5II upright confocal/multi-photon microscope (*Leica Microsystems, GmbH, Wetzlar, Germany)* (λ_Ex_: 880 nm, λ_Em_: 500-600 nm). The laser source consisted of a visible Argon laser (power = 29%) with a multiphoton gain of 31% and multiphoton offset of 63%. Emitted photons were detected on a PMT NDD1 detector at high voltage (786 V). In parallel to p3, the nucleus of cells was simultaneously detected using the nucleotide binding dye DAPI via confocal microscopy (λ_Ex_: 405nm, λ_Em_: 400-500 nm). DAPI was detected using a 405 nm UV laser diode with emitted photons detected on a HyD 1 detector at 82% gain. Cells were imaged at a bidirectional scan speed of 600 Hz using a 40x oil immersion lens via Type F Immersion Liquid (NA= 1.25 and RI = 1.52). Collected images were line averaged (level = 4) and had a resolution of 8 bits (1024 x 1024 pixels). Final images from the two channels were merged and processed on the LEICA LAS X Imaging Software and exported with scale bar. H2S levels were subsequently quantified using open-source ImageJ (https://imagej.nih.gov/ij/index.html) by measuring the integrated density per unit area for each cell in the p3 channel. Position of cells were confirmed using the DAPI channel.

### Cell Migration Assay

Assay was initiated by scratching with a 10 µL pipette tip along the bottom and center of a well in a 6-well plate. A straight edge was used to ensure the scratch was a straight as possible. After scratching, cells were rinsed twice with 1x PBS to remove detached cells, and siRNA. Next, 2 mL of DMEM (1% FBS, 1% P/S) was added to cells with or without H_2_S treatment using 2-fold dilutions. Low concentrations (≤100 µM) of H2S were delivered via NaHS while higher concentrations via Na_2_S (≥200µM). Immediately after treatment, cells were transferred to a LEICA inverted microscope equipped for live cell imaging (37°C temperature, 5% CO_2_ control) in the Lerner Research Institute Imaging Core. Images were acquired using a Leica DMI6000 inverted microscope (Leica Microsystems, GmbH, Wetzlar, Germany) equipped with a Hamamatsu Orca Flash4 camera (Hamamatsu Photonics, Shizuoka, Japan). Cells were imaged using phase contrast at 10x air phase magnification (NA = 0.4, RI = 1), exposure time = 24.49 s, intensity = 50%, and 2×2 CCD camera binning at 8-bit resolution for 24 hours with 30 min time steps. Five positions were imaged per well. Images were processed via the LEICA LAS X Imaging software and exported as.tiff file with scale bar and elapsed time.

### Gap quantification with ImageJ

Gap was quantified using open-source ImageJ software downloaded from the NIH (https://imagej.nih.gov/ij/index.html). Lines were drawn around the gap for time points 0, 6, 12, 18, and 24 hours to calculate the area. Gap was quantified as “Area” in ImageJ and represented as a percentage normalized to 0 hr for each position (Eq.1).

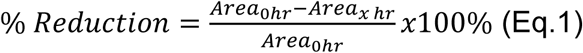

### Gap quantification with CellProfiler

Gap quantification was automated using open-source CellProfiler software downloaded from https://cellprofiler.org. The gap was determined by subtracting the total image area with the total cell area. To calculate the total cell area, images were processed using the following steps. An image was first cropped to remove the elapsed time and scale bar. The cropped image was then converted into grayscale, where edge detection (canning algorithm with absolute threshold = 0.5 and automatic sigma detection) was used to increase edge contrast. Circular objects (indicative of the cells) with a size of 40 were subsequently enhanced. Finally, a smoothing algorithm (gaussian filter with artifact diameter = 100) was used to merge enhanced features into distinct groups. Ideally, there should be two main groups after this step representing the two migrating fronts of the cells. After image processing, the total cell area was calculated by identifying all features with size between 10 and 40 (roughly the size of the cells) using a global, two-class Otsu thresholding algorithm. The gap area was then determined by subtracting this cell area from the total image area. Gap area was reported as both gap area (in pixels) and as a percentage of image area. Final segmented images were exported and saved in the designated storage location. The CellProfiler pipeline file used in this study is provided as part of the supporting information.

### Bismuth Sulphide Detection Assay

20 mM of bismuth (III) acetate (Sigma Cat# 401587) was dissolved in a solution of 15% glacial acetate acid (Sigma Cat# 695092). Deionized water (Milli-Q IQ7000 Ultrapure Water System) was used for dilution. Total volume for each batch consisted of 100 mL. Blotting papers cut to size of 96-well plate were soaked in the prepared bismuth acetate solution for 20 minutes at room temperature. The soaked papers were then dried overnight in a vacuum chamber. Dried papers were either immediately used or stored under vacuum. For the actual detection, hydrogen sulfide was detected as bismuth (III) sulfide based on a previously published protocol (29).

### Statistical Analysis

All measurements are expressed as a box plot showing the arithmetic mean, maximum and minimum value, along with individual data points. The sample size (*n)* for all experiments was n ≥ 5. Where relevant, hypothesis testing was conducted between groups either using Student’s t-test or One-Way ANOVA. Statistics were conducted on GraphPad Prism9. A p-value of *< 0.05* was determined to be statistically significant, if not otherwise indicated. *p<0.05, **p<0.01, ***p<0.001, ****p<0.0001.

## Acknowledgements and Funding Sources

The authors acknowledge financial support from the Lerner Research Institute, Center for Transformative Nanomedicine, Puzzitelio Family Foundation. A.C. acknowledges support from the National Science Foundation Graduate Research Fellowship. The authors would like to thank Dr Nora Singer from the Cleveland Metrohealth for providing the HaCaT cell line, and Dr Ajay Zalavadia from Cleveland Clinic Research Imaging Core for microscope training and guidance.

## Author Contributions

A.C. and V.K. conceived the idea, designed the experiments, and wrote the manuscript.

A.C. performed the cell migration experiments.

## Data Availability

The authors declare that all data supporting the findings of this study are available within the manuscript and supplementary information.

