## Supporting Figures for "Hydrogen Sulfide Suppresses Keratinocyte Migration in Scratch Assay"

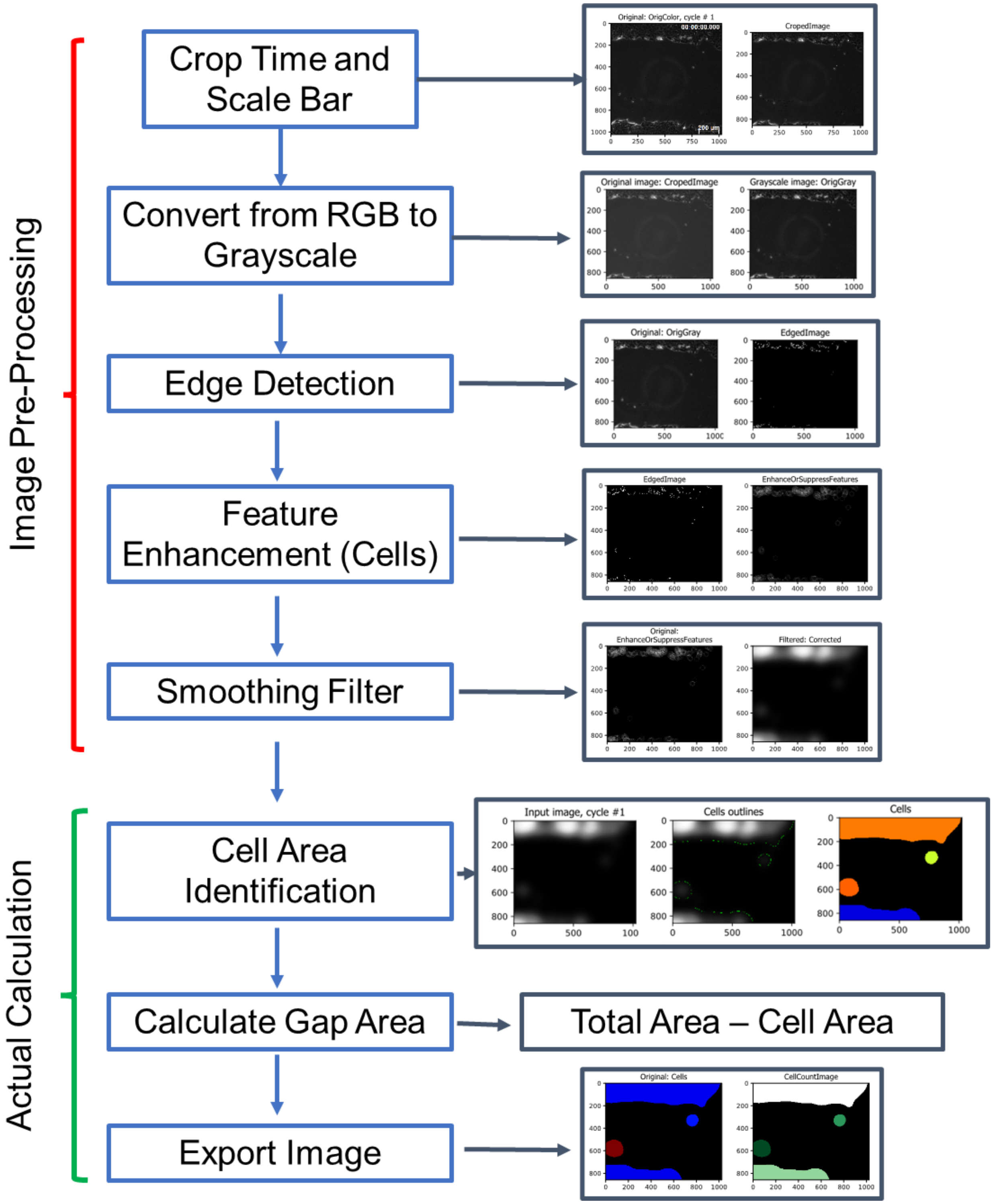


**Supporting Figure 1.** Walkthrough of the pipeline for the CellProfiler algorithm developed on a single image (HaCaT Keratinocytes: Control, Trial 1 Position 1).

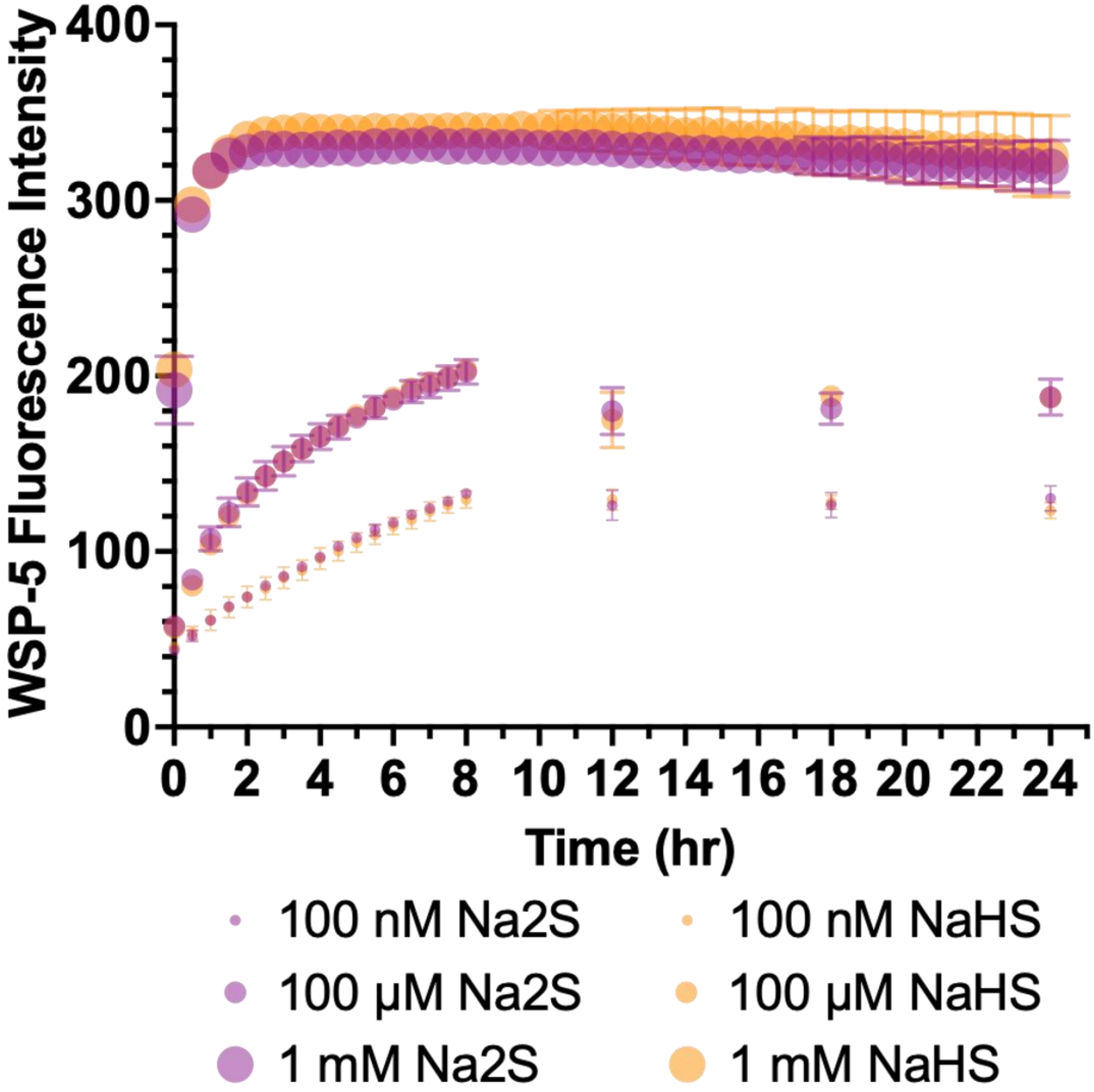


**Supporting Figure 2**. Release comparison for NaHS and $Na_2S$ as detected using WSP-5 dye.

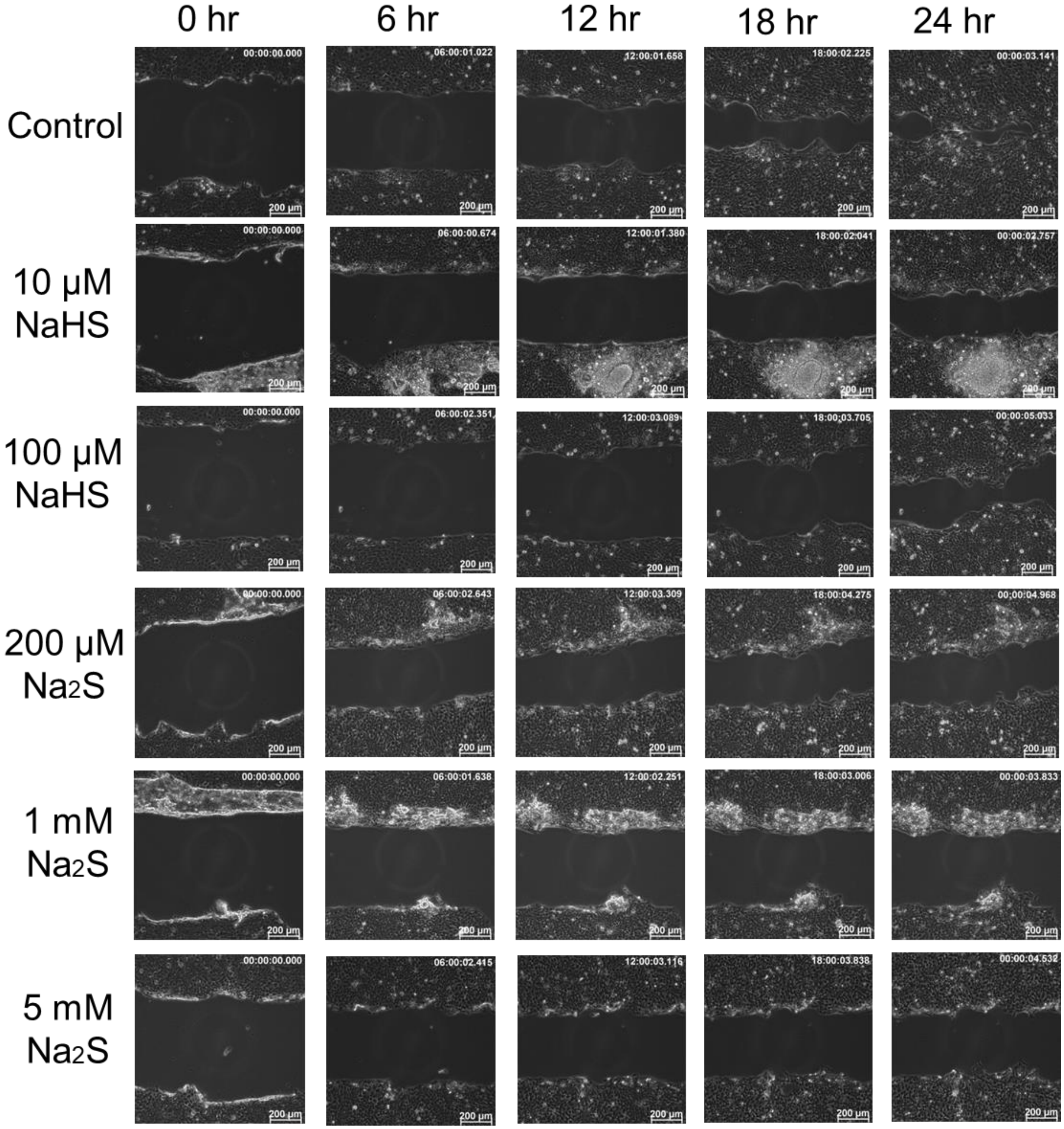


**Supporting Figure 3**. Snapshot images of HaCaT migration for all exogenous donor concentrations are shown at 0, 6, 12, 18, and 24 hours.

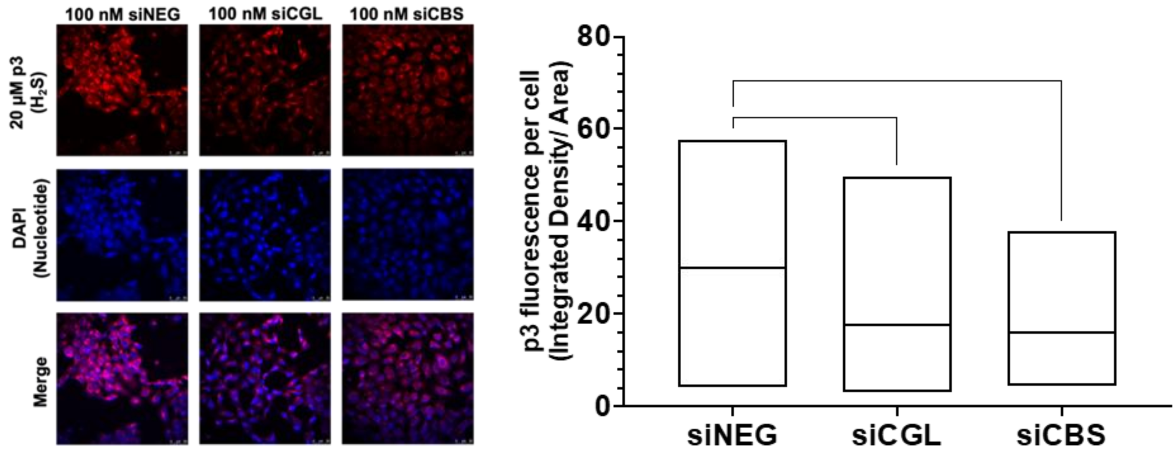


**Supporting Figure 4**. Investigation of endogenous $H_2S$ production in HaCaT keratinocytes (passage 60 ± 2) using mRNA silencing of $H_2S$-producing enzymes cystathionine-γ-lyase (CGL) or cystathionine-β-synthase (CBS) for development of cell migration scratch assay. Lipofection consisted of 100 nM siRNA complexed in Lipofectamine RNAiMax. SiRNA silencing was evaluated in cells using p3 multi-photon fluorescence probe (λex = 880 nm, λem = 500-600 nm) to detect $H_2S$ levels. $H_2S$ levels were quantified as intensity density per area using ImageJ. (n = 70-128 cells).

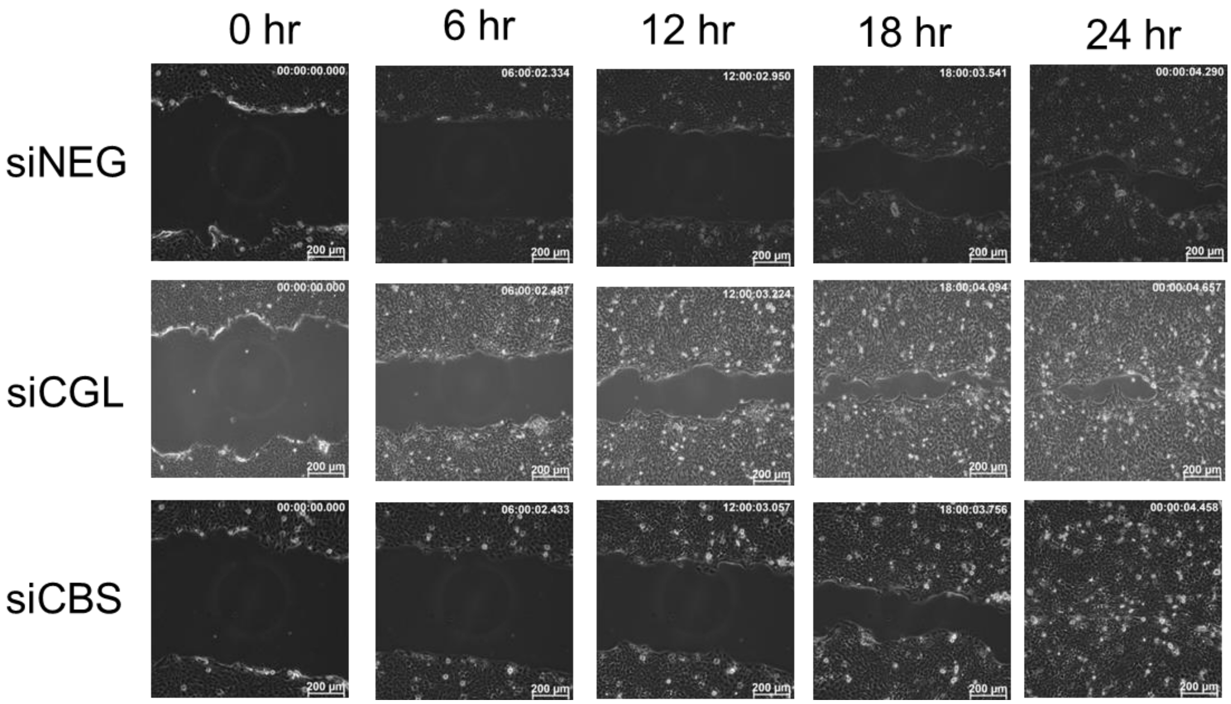


**Supporting Figure 5**. Snapshot images of HaCaT migration for all endogenous donor concentrations are shown at 0, 6, 12, 18, and 24 hours.
